# Urbanization Affects Nest Material Composition and Use of Anthropogenic Materials in Blue Tit Nests (*Cyanistes caeruleus)*

**DOI:** 10.1101/2024.04.25.591121

**Authors:** Joseph Roy, Davide M. Dominoni

## Abstract

Avian nests are complex structures that protect eggs and nestlings. Nest-building behaviour varies between species and habitats, and recent work has highlighted that in areas with high human activity and low availability of natural nest material, birds may use anthropogenic material to construct nests. However, we still know relatively little about how nest composition is affected by human presence, including along urban gradients. Here we present the results of a study where we examined how nest composition differed between an urban and a forest population of blue tits, and the impact that variation in nest composition had on reproductive success. We found a significant decrease in the amount of natural materials and an increase in anthropogenic materials in urban compared to forest nests. Earlier urban nests showed a higher amount of anthropogenic materials compared to late urban nests. The amount of moss incorporated in a nest was the strongest predictor of fledgling success, suggesting that the use of anthropogenic material by urban birds might be a maladaptation, and/or that urban birds are constrained in the amount of moss they can use to build their nests. Future studies should aim at quantifying the availability of natural materials to distinguish between these two hypotheses.

## Introduction

The process of urbanization can have various impacts on ecosystems, such as the elimination of natural vegetation and the construction of artificial and impermeable structures (Sepp et al., 2018), the introduction of pollutants, including light and chemicals (Dominoni et al., 2013), and the increasing levels of human disturbance (Sol et al., 2013). There may be relaxation in ecologically significant factors such as predation pressure (Eötvös et al., 2018), food availability (such as an increase in lower quality food) (Marzluff, 2001) and temperature (due to the urban heat island effect) (Puppim de Oliveira et al., 2014). Whether a species can thrive in an urban environment depends on how it perceives the changes brought about by its new habitat. Some animals can adapt and take advantage of the opportunities presented become exploiters, and adapt and survive. Unfortunately, some animals may experience a population decline and avoid the area altogether. (Kark et al., 2007; González-Lagos & Quesada, 2017; Corsini et al., 2021). The urban adapters might have to change their life histories to adapt to the conditions and they will have to utilize the available resources. Urban dwellers adapt and adjust their life histories to cope with changes in their environment. Changes in circadian rhythm, foraging behaviour, communication strategies, vigilance behaviour and changes in predator response are some examples (Marzluff, 2001; Sol et al., 2013; Tätte et al., 2019).

In birds, nest building behaviour is a crucial step of the breeding phase. Several studies have shown that nest building can change in urban areas. These changes can involve using less natural materials, altering the structure of the nest, and incorporating man-made materials (Antczak et al., 2010; Votier et al., 2011)(Jagiello et al., 2019). The term “anthropogenic nest material” refers to items not naturally found in nature but created by human actions. Materials such as plastic, string, and coir are examples of anthropogenic nest materials (Briggs et al., 2023). The incorporation of human-derived materials is shown to have a positive correlation with the human footprint index (Z. Jagiello et al., 2019). In urban environments, the availability of natural nest materials is often reduced, leading birds to use debris as a replacement (Wang et al., 2009). Synanthropic birds (those that thrive in environments modified by human activity, such as cities) may incorporate more debris into their nests than non-synanthropic birds (those that do not adapt well to human-modified environments) (Z. Jagiello et al., 2019). This raises the question of why birds use anthropogenic materials and how it affects their life histories.

Jagiello et al., 2022 provided three major hypotheses that explain anthropogenic material use in bird nests. The availability hypothesis states that the availability of anthropogenic materials and the absence of natural materials causes the increased use of the former. A study that looked at the preference for nest material in Pied Flycatchers found that when available, the birds preferred natural material over human-derived material (Briggs et al., 2023). The age hypothesis proposes that the age of parent birds and their experience is connected to the use of anthropogenic material, where old and experienced birds use it often as an extended sexual signal (Sergio et al., 2011). Finally, the functional hypothesis states that the functional advantage of the anthropogenic materials leads to their use in nests.

Several advantages have been reported for the use of human-derived materials in bird nests. Undoubtedly, plastic strings have their advantages, as they play a crucial role in enhancing the nest structure, which is a key factor in safeguarding the eggs and fledglings from harsh weather elements like the wind (Deeming, 2004). Manipulating these materials during nest building could be significantly easier than using natural alternatives (Esquivel et al., 2020). Shrikes, for instance, are known for impaling various objects and using them for construction. Another benefit of having many man-made materials readily available in the area is that it can lower the effort and time spent searching for nesting materials, leading to faster nest building (Antczak et al., 2010). Cigarette butts are found to have a repelling effect comparable to green materials, which could decrease the number of ectoparasites found in city bird nests (Suárez-Rodríguez et al., 2013). A study in Black kites showed that older birds showed more tendency to incorporate plastic material in their nests up to an age and they stopped using plastic after senescence. This suggests that the use of plastic might be used as a signal of sexual quality (Sergio et al., 2011).

There are also several potential negative impacts of anthropogenic nest materials. Modifications of nests might alter their structural and functional properties including integrity, camouflage and thermoregulation (Mainwaring et al., 2014; Lopes et al., 2020). Suárez-Rodríguez et al., 2013 looked at the effects of cigarette butts and its effects on breeding success and found that even though butts have ectoparasite-repellent effects, the number of cigarette butts in the nest was positively associated with the level of female genotoxic damage in house finches during breeding. The study also found that the negative effects of cigarette butts on the breeding success of urban birds outweigh any potential benefits. Entanglement of chicks in the plastic debris is reported in a study of a colonial seabird Northern Gannet (Morus bassanus) (Votier et al., 2011). Plastic waste can also cause chemical harm through the release of toxins, leachates, and non-degraded persistent organic pollutants which can enter food webs via a process known as trophic transfer (Teuten et al., 2009). As mentioned, camouflage is an important characteristic to prevent nest predation. Predation is a major cause of nest failure and several studies have shown the effects of artificial nesting materials on predation rates (Corrales-Moya et al., 2023). In their 2010 study, Antczak et al. noted in their study of great grey shrikes (*Lanius excubitor*), two nests that had entire white plastic strings, made them easier to be detected by predators. In a study on clay-coloured thrush (*Turdus grayi*), the proportion of the outer layer of the nest covered by artificial materials was negatively correlated with the nest’s daily survival rate, indicating that the more exposed artificial material on the nest, the higher the chance of it being predated (Corrales-Moya et al., 2023). The study analysed whether the internal or external dimensions of the nest had any impact on the nest survival rate. The findings revealed that variations in the nest dimensions did not affect predation events while the only factor that had an effect was exposed anthropogenic material. Although plastic is considered a thermal insulator, in hotter climates this might act negatively causing thermal risk for the chicks by increasing the temperature in the nests. In American crow (*Corvus brachyrhynchos)*; breeding success was significantly low in nests with higher amounts of plastic. A major reason for the reduction was the entanglement of the chicks, but even in un-entangled nests, the success was low owing to the negative effects of plastic in nests (Townsend & Barker, 2014). A similar case of strangling was reported in Great Gray Shrikes (*Lanius excubitor*) and it is suggested that both adults and the chicks are prone to strangling (Antczak et al., 2010).

Although several studies have looked into the effect of anthropogenic materials on breeding success, most of these have focused on marine habitats and sea birds and only few on passerines (Jagiello et al., 2019). A study on blue tits and great tits found that the presence of anthropogenic materials was negatively correlated to breeding success in blue tits but not in great tits (Jagiello et al., 2022). In the current study we aim to expand on these previous findings and understand 1. the effect of urbanization on nest composition, and 2. the relationship between nest composition and breeding success. To this end we collected nests and breeding data on blue tits (*Cyanistes caeruleus*) breeding at nine sites between the city of Glasgow and a nearby forest in the Loch Lomond and Trossachs National park, UK.

## Methods

### Monitoring of Breeding Success

The study was carried out at the urban-rural gradient at the University of Glasgow, Scotland, using nine sites spanning 35 miles transect from the city centre of Glasgow (55.8668° N, 4.2500° W) to the Ross woods at the east banks of Loch Lomond (56°7’44”N 4°36’46”W). Over the past few decades, approximately 500 Schwegler Woodcrete nest boxes with 32mm entrance diameter have been placed along this gradient and monitored for hole-nesting birds (Reid, 2020). From April 1st to June 15th, 2023, breeding success was monitored across all sites. Once the birds started nesting, all the boxes were checked once a week to assess the progress of the nest building and incubation. Close to the expected hatching time we started to check the active nest every other day until chicks were found. Once the chicks were 10 days old, they were ringed by BTO licenced ringers, and morphological measures (wing length, tarsus length and weight) were collected. Once the chicks were fledged the data for breeding success was collected by counting the number of unhatched eggs and dead chicks in the nest boxes and the nests were collected in plastic zip-lock bags.

### Nest dissection

Once the nests were collected, they were frozen for at least 12 hours to reduce the activity of arthropods and they were kept frozen until the time of dissection. Before dissection, the nests were dried in a heat chamber at a temperature of 50 degrees Celsius for at least 12 hours to prevent moisture in the nests. Once dried the nests were dissected and each category was measured up to an accuracy of 0.001g. The materials were grouped into four categories: moss and grass (natural materials), feathers, wool and hair (naturally derived), and anthropogenic material (together they will be referred to as nest materials from now on). As these materials are intertwined intricately, maximum efforts were taken to separate the materials into each class.

### Environmental Variables

Habitat was defined selected as a categorical variable with two levels (urban and rural) to represent that the nests were collected from the far extremes of the urban-rural gradient. We also extracted the impervious surface area of around 100m (IS100 from now on) of the nest box from available land cover maps (Imperviousness — Copernicus Land Monitoring Service, 2018). This variable was used by Z. Jagiello et al., 2022 in a similar study that looked at anthropogenic materials. The impervious surface area was selected as a proxy for the amount of urbanisation.

### Statistical Analysis

The statistical analysis was performed using R version 4.3.3 (2024-02-29 ucrt)-- “Angel Food Cake”. The analysis was conducted in two parts. In each of them, model selection was conducted via likelihood ratio tests and the jPlot package was utilized to visualize model predictions.

#### Nest Materials

To assess environmental effects on nest composition, we used Linear Mixed-effect Models (GLMMs) with each nest material type as response variable, and site as a random variable. Two sets of models were run for each nest material type, with the following fixed effects: 1) habitat (urban and rural), first egg date and their interaction; 2) IS100, first egg date and their interaction. We did so because habitat was collinear with impervious surface area, but we still wanted to distinguish between the effect of habitat per se versus the effect of a specific environmental variable that might better function as a proxy of human activity and thereby the use of anthropogenic materials in nests. We perform model selection only on interactions. That is, if an interaction was not significant we removed it and considered the final model the one with all linear terms. We visualized model predictions using the plot_model() function in the sjPlot package.

#### Breeding Success

From the breeding dataset, clutch size, hatching success (number of hatched eggs/clutch size) and breeding success (number of fledglings/number of hatched eggs) were extracted. Clutch size was modelled with Poisson GLMMs while hatching and fledging success were modelled with Binomial GLMMs. In all models, nest material type and first egg date were included as fixed effects, and site as a random effect. We visualized model predictions using the plot_model() function in the sjPlot package.

## Results

### Nest Materials

We focused on five nest material variables: 1. total weight, 2. anthropogenic materials, 3. moss + grass, 4. Feathers, 5. wool + hair. For the total weight, the values span from 7.45 to 37.01g, with a median of 19.45 and a mean of approximately 20.35g. Urban nests had the highest nest weight and had more variability than the rural nests. Anthropogenic material ranged from 0.00 to 7.89g. The median was just 0.008g, pointing to a skewed distribution with a mean of around 0.45g. Anthropogenic materials were near zero with a mean of 0.02 and a median of 0.00 for rural nests, while the mean and median for urban nests were 0.92 and 0.40, respectively. In the rural habitat, 45.45% of nests contained anthropogenic materials while the urban nests had 90.32%. Combined, anthropogenic material was present in 71.87% of the nests (Figure 3). Moss and grass values ranged from 4.83 to 28.98g, with a mean of 14.38g and a median slightly below the mean at 14.10g. Rural nests had more moss and grass with a mean value of 15.4, while urban nests had a mean of 13.3. In the case of feathers, the values spanned from 0.035 to 7.02g. The mean was around 1.06g with a median of 0.81g. Feather weight was higher in the urban nests with a mean of 1.20g, while the rural nests showed a mean of 0.94g of feathers. Finally, wool and hair values ranged from 0.00 to 11.44g, showing a mean of 2.29g and a median of 1.41g. Wool and hair weight was higher in the urban habitat with a mean of 3.41g, while the rural habitat had a mean of 1.25g. Mean, median and SD values for all these categories are available in Table 2.

**Figure 1.**
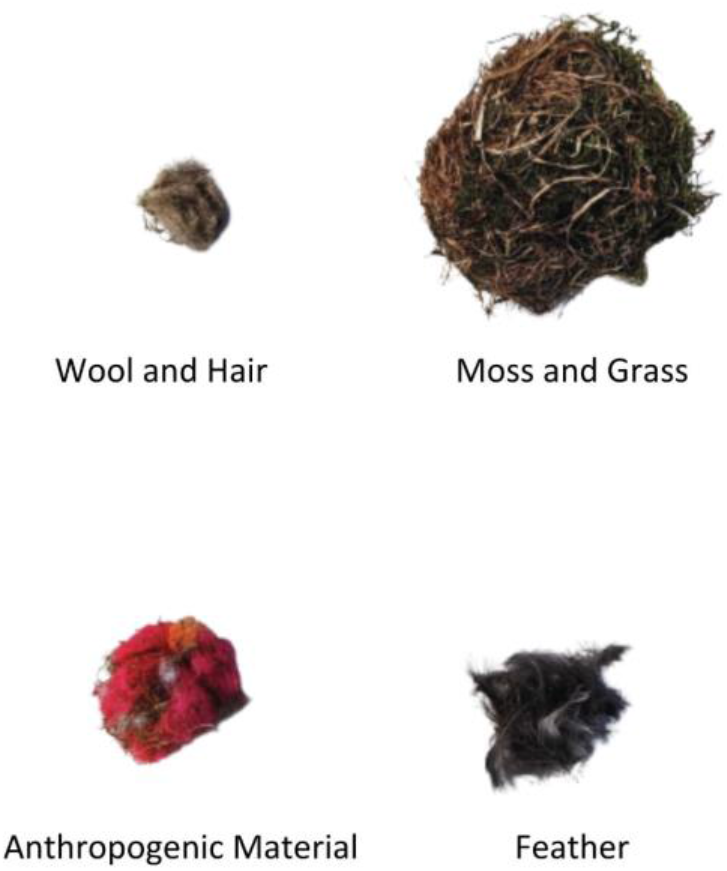
Picture showing different categories of nest materials.

**Figure 2.**
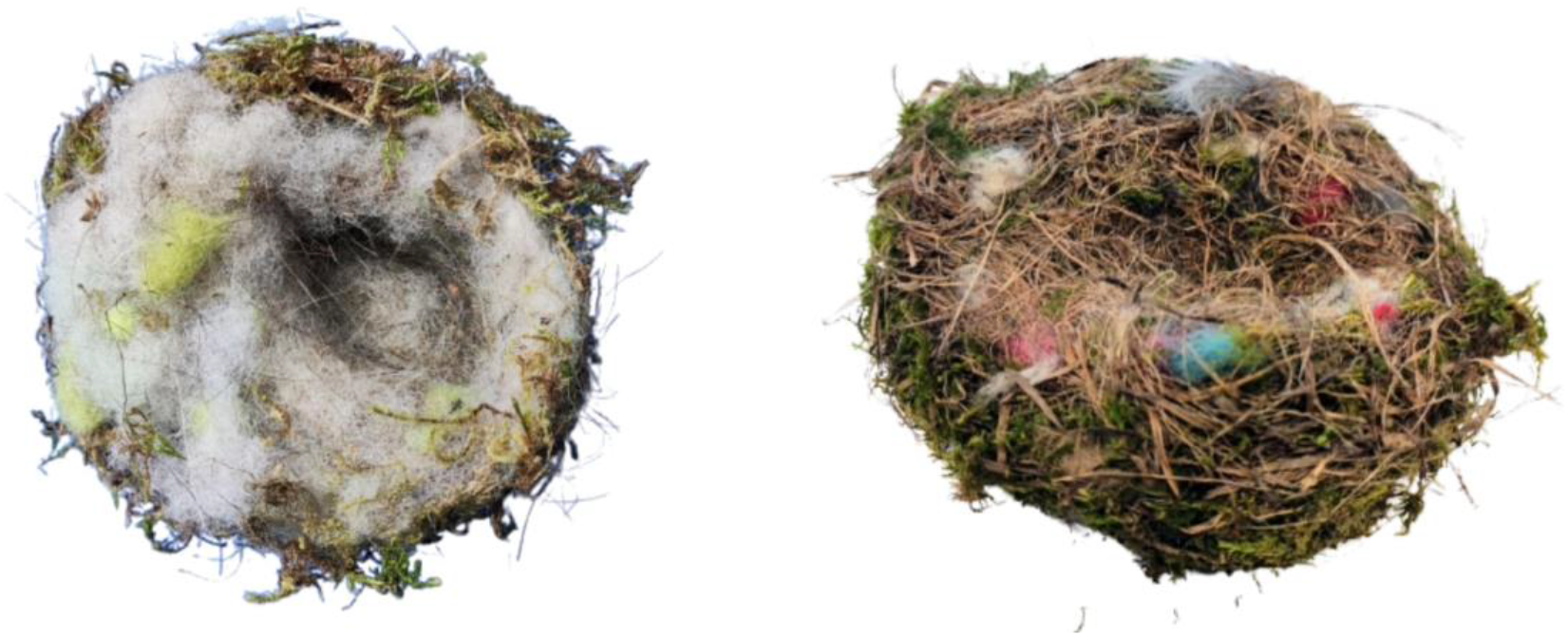
Images of nests with anthropogenic materials. On the left-a nest with the whole cup made of human-derived fibres. The right image shows a multicoloured nest.

**Figure 3.**
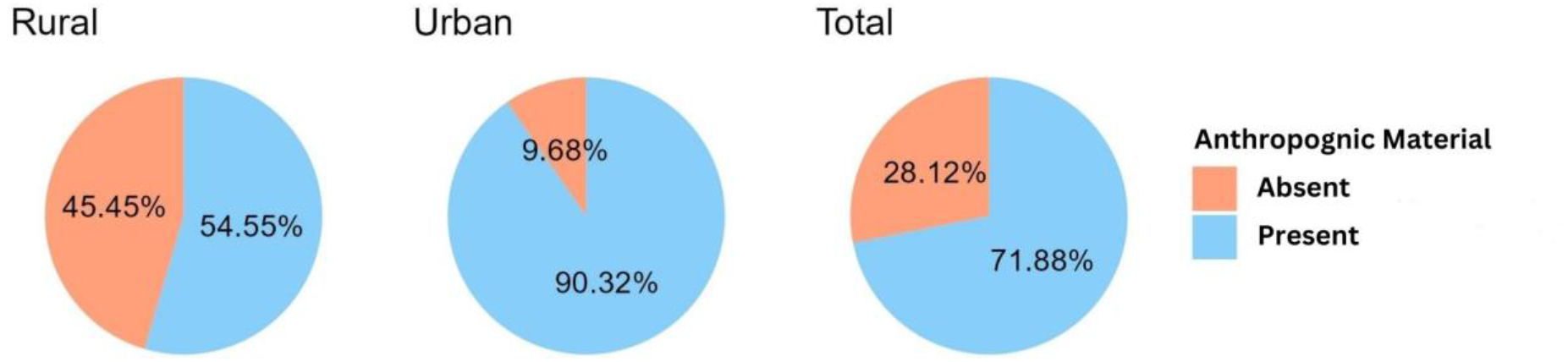
Pie charts showing the presence of anthropogenic materials in the nests. There was anthropogenic material in half of rural nests while 90% of urban nests had anthropogenic materials in them. In total three-quarters of the nests had anthropogenic materials.

**Figure 4.**
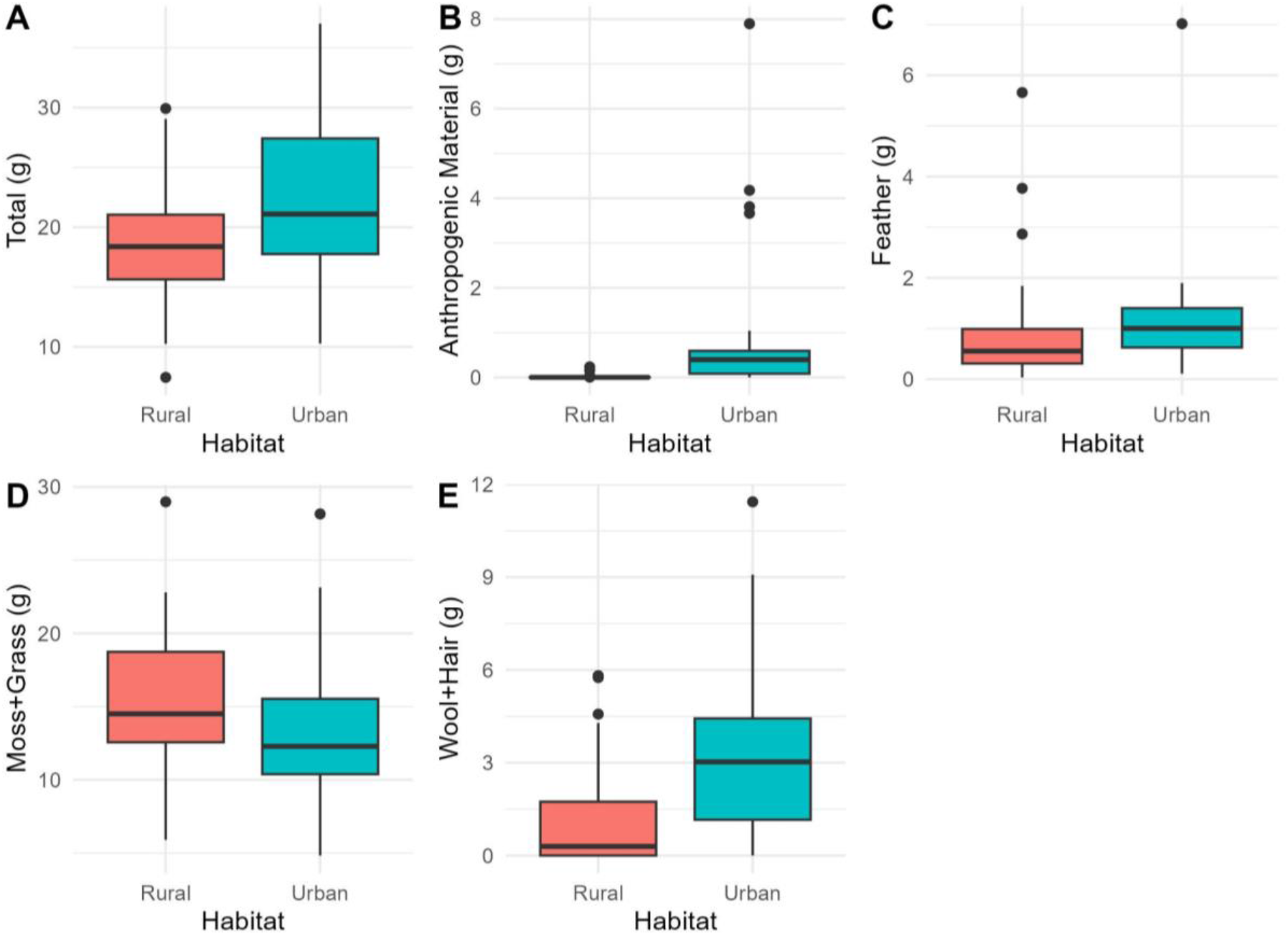
Box plot representation of Nest Materials from urban-rural habitats.[A] Shows the total weight of nests, Rural habitat has fewer heavy nests while their weights are mostly around 20g while urban nest weights are variable.

**Figure 5.**
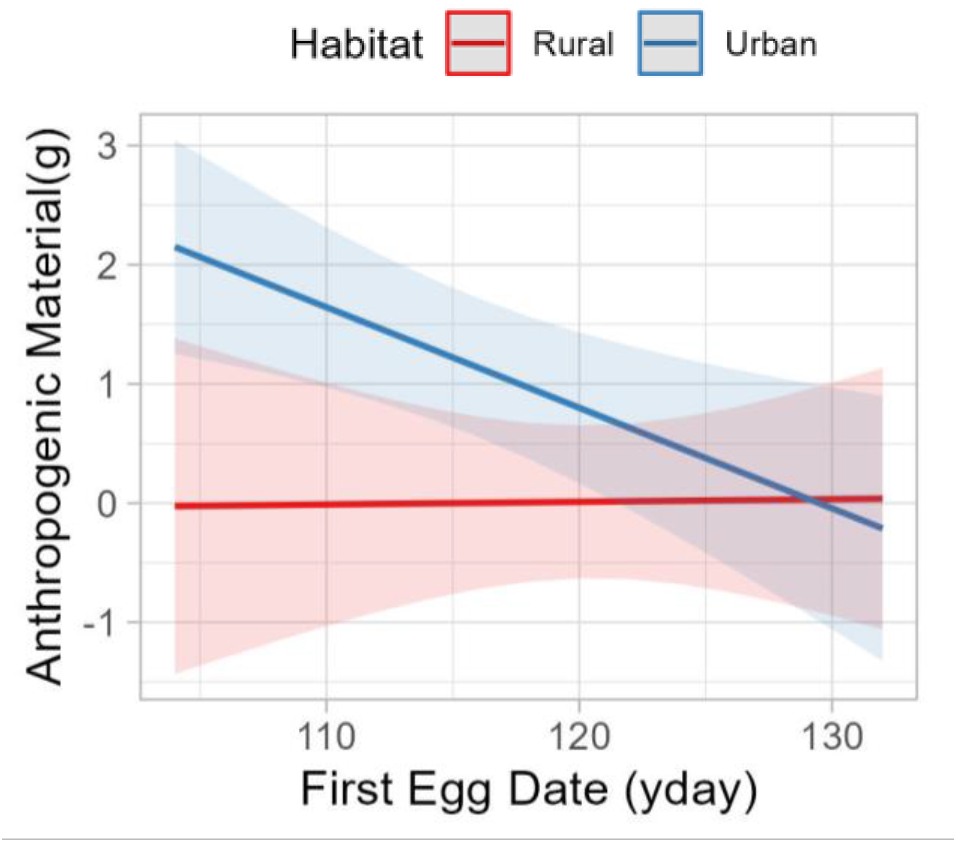
Predictions of a model explaining variation in anthropogenic nest material as a function of habitat and first egg date. The solid red line represents fitted linear regression and the shaded area represents 95% confidence intervals.

In the models with the nest materials as response variables and the interaction between first egg date and habitat (urban/rural) as fixed explanatory variables, models showed no significant effect on except for the models focused on anthropogenic materials. Here, the interaction between habitat and first egg date showed a marginal significant effect with a negative slope for the urban environment, suggesting that when building nests in urban habitats earlier in the spring the likelihood of incorporating anthropogenic material was higher (estimate= -0.09, p-value= 0.07). The linear term of habitat was significant and suggested that urban nests incorporated more anthropogenic material than rural nests (estimate = 12, p-value= 0.05). Summary tables of these results can be found in the supplementary material (table 1-5).

**Table 1.**
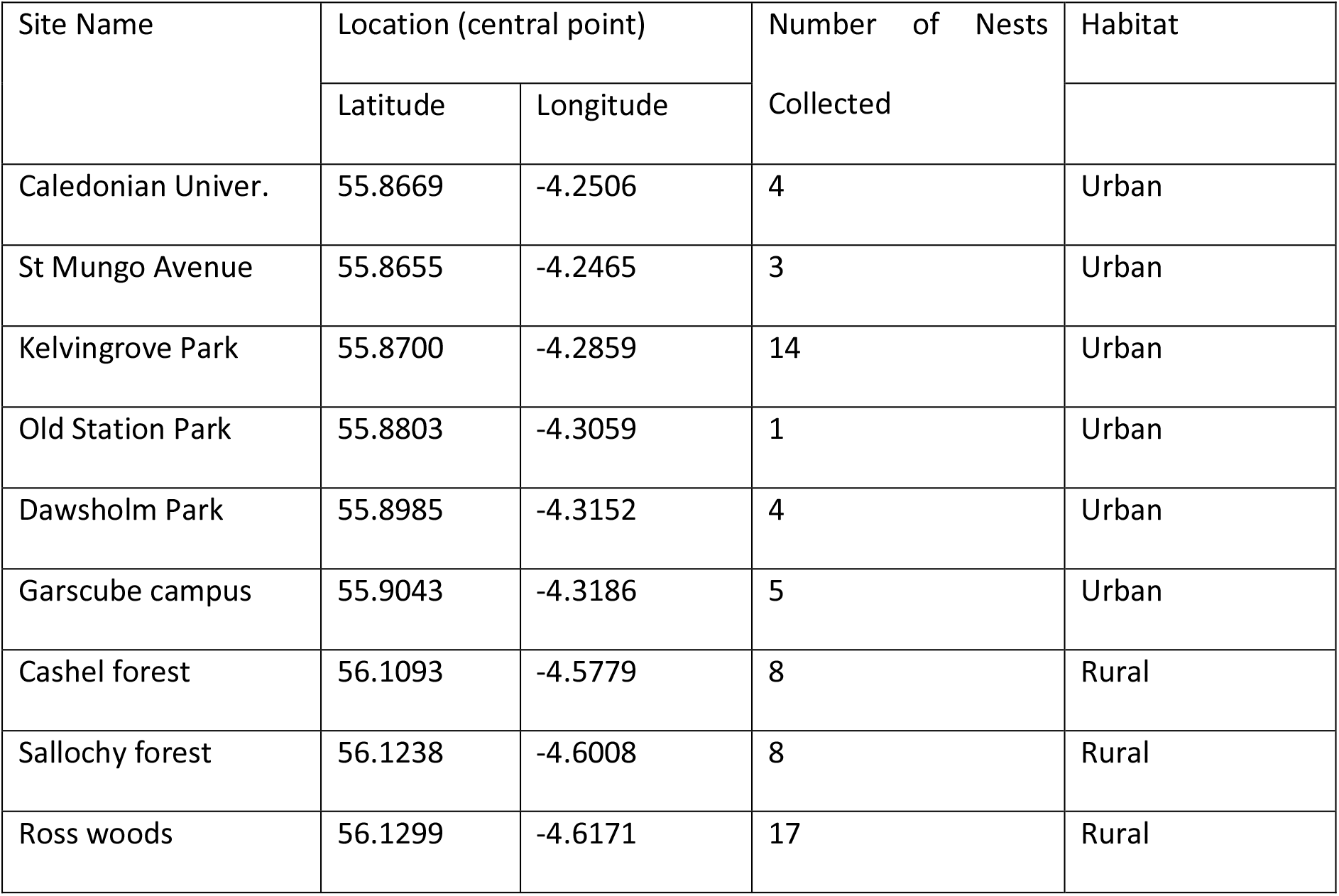
Table showing the sites, their coordinates, number of nests collected from each sites and their respective habitats. From six urban sites and three rural sites a total of 64 nests were collected.

**Table 2.** Table representing mean, median and standard deviation of the weight of each nesting material type found in blue tit nests.

| Characteristic | Rural, N = 33<sup>1</sup> | Urban, N = 31 |
| --- | --- | --- |
| Total | 18, (18), (5) | 22, (21), (7) |
| Anthro | 0.02, (0.00), (0.05) | 0.92, (0.40), (1.70) |
| Moss+ Grass | 15.4, (14.5), (5.1) | 13.3, (12.3), (4.9) |
| Feather | 0.94, (0.55), (1.16) | 1.20, (1.00), (1.20) |
| Wool+ Hair | 1.25, (0.30), (1.80) | 3.41, (3.03), (3.05) |

<sup>1</sup>Mean, (Median), (SD)

We then assessed the effect of a continuous rather than a dichotomous variable representation urbanisation, the amount of impervious surface (IS100) around each nest location, on the mass of each nest material type. In all models, the interaction was not significant and hence only linear terms were retained. For anthropogenic materials, the first egg date showed a statistically significant negative trend (estimate= - 0.42, p-value= 0.01). In the case of Moss and Grass, IS100 showed a significant negative trend with the increase in IS100 (estimate= -2, p-value= <0.01). The predictive graphs are added as figure 6.

**Figure 6.**
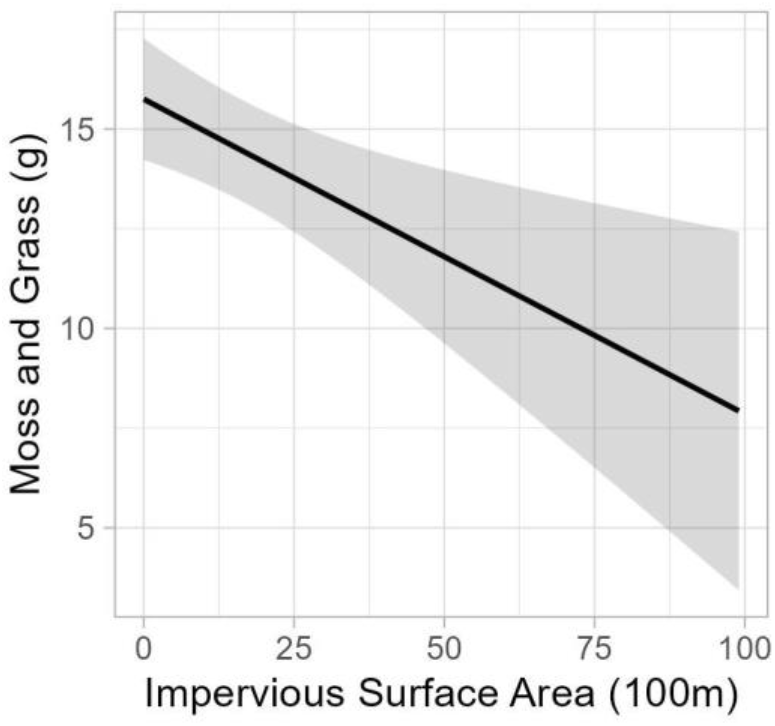
The solid red line represents fitted linear regression and the shaded area represents 95% confidence intervals, the predictive graph of the change in the weight of moss with an increase in impervious surface area in 100m. There is a statistically significant reduction in the weight of moss with the increase in the amount of impervious surface area.

### Breeding Success

The models with nest materials and first egg date showed no statistically significant results for clutch size and hatching success. However, for fledgling success, model selection indicated that the best model was the one where only moss and grass were retained. This variable showed a positive effect on fledgling success with a marginally significant p-value (estimate= 0.26, p-value= 0.06). The predictive graph is shown in Figure 7 and the detailed results and model selection tables are added in the supplementary material (table 11-13).

**Figure 7.**
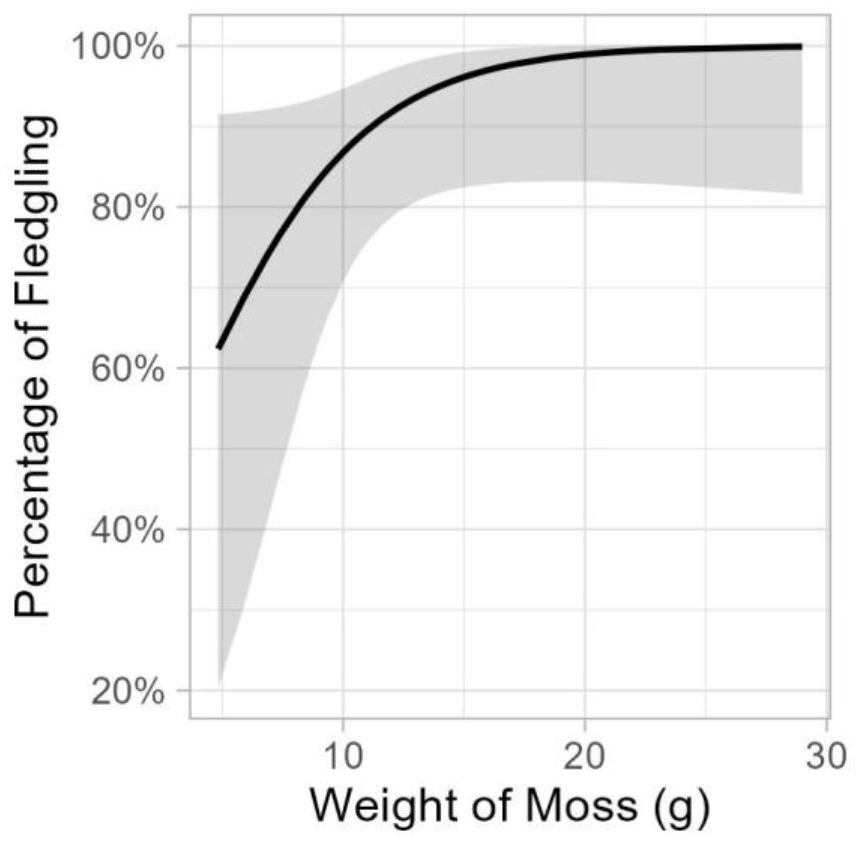
Predictive graphs on fledgling success and the variation in the weight of moss. The solid red line represents fitted linear regression and the shaded area represents 95% confidence intervals. Fledging success is increased with the increase in the weight of Moss and Grass.

## Discussion

The findings from the present study shed light on the influence of nest materials and urbanisation on bird populations and their breeding success. We found that the rural nests were significantly heavier and had more natural materials, particularly moss and grass, compared to the urban nests, while the latter contained significantly more anthropogenic material. The amount of moss and grass was a strong predictor of fledging success: nests that contained more moss and grass fledged more young.

Looking at the individual materials, the combination of moss and grass (natural materials) was the only material that showed a significant reduction with an increase in impervious surface area. Total weight and wool also showed a reduction with IS100 but did not show significance. Moss and grass emerge as significant predictor of fledgling success which indicates the essential role of natural materials in nest construction, by providing better insulation or structural support. In these models, anthropogenic materials did not show a significant effect on breeding success when it was assessed along with other materials. This underlines the importance of natural materials in the breeding outcome.

Even though anthropogenic material was not significant, there was a notable increase in the amount of anthropogenic materials in the urban nests. Also, it is to be noted that 72.8% of all the nests had the presence of anthropogenic material in them. We have noted that in urban nests, anthropogenic material was mostly used in cups to provide structural support instead of grass (J.Roy pers obs). The reduced availability of grass might be the reason for birds to use anthropogenic materials in their nests. The anthropogenic materials were mostly fibres from tennis balls, fabrics and plastic threads which serve the purpose of providing structural integrity. The selective picking of nesting material was demonstrated by Briggs et al., 2023 and it was observed in several previous studies. Even though urbanization can be considered as a proxy for the reduction of materials like moss and grass, it can only be confirmed if these materials are selected out of choice or out of need. The amount of anthropogenic material was predicted to increase the number of unhatched eggs Jagiello et al., 2022; James Reynolds et al., 2019. It is to be noted that this study mostly focused on anthropogenic material while in our results moss had the most effect on breeding success. James Reynolds et al., 2019 suggest city-specific differences in nesting behaviours, the changes we observed can be due to differences in the availability of materials to the different micro-habitat which is offered by nesting locations. Suárez-Rodríguez et al., 2013, in a similar study, reported cigarette butts in blue tit nests as an insect repellent, no nests in our sample had cigarette butts in them.

Several theories have attempted to explain the reasons why birds incorporate anthropogenic materials in their nests. When provided with natural materials, birds showed a preference for natural materials over anthropogenic nest materials (Lee et al., 2015). However, in our study, the three rural sites were oak woodlands with ample amounts of natural materials and anthropogenic materials are scarce and are mostly the trash from occasional tourists (J.Roy pers obs). Still, 54.5% of rural nests had anthropogenic materials in them (*Figure* ) even though the weights were considerably lower than in urban nests as shown in *Table* . The presence of anthropogenic materials in the rural nests, where natural materials readily available, might show a preferential selection towards anthropogenic material in rural habitats as observed by Briggs et al., 2023. As previous studies have suggested, birds tend to use anthropogenic materials as an extended sexual phenotype. In the case of rural habitats, the anthropogenic materials were mostly used along with feathers which might play a decorative role in the nests. Such a preferential selection of anthropogenic materials could show a selective preference and potentially using anthropogenic materials as an extended sexual phenotype. Feathers in nests are associated with increased thermal regulation and resistance against microbial and ectoparasite infections (Järvinen & Brommer, 2020). On these accounts, Jagiello et al., 2022 suggested that the chicks from feather-rich nests have a higher chance of recruitment to the breeding population as the number of feathers shows female reproductive quality. As presence of anthropogenic materials in nests might affect mate choice which can in turn affect breeding success.

In urban habitats, anthropogenic materials were often incorporated into the structural parts of the nests instead of grass which could mean that birds are using these materials for stabilizing the nest structure. In urban habitats, 90.3% of nests had the presence of anthropogenic materials, also the weight of anthropogenic materials was significantly high in comparison to the rural habitats. The reason for urban birds to use anthropogenic nests cannot be concluded without quantifying the availability of natural nesting material in urban habitats. Although there is a marked increase in the amount of anthropogenic materials, breeding success was mostly driven by the total weight of the natural materials (moss and grass). Also, the earlier nests had a significantly higher weight of anthropogenic material in the city while in the rural areas, the amount of anthropogenic materials stayed low throughout. Earlier laying date is considered to show parent quality and our study has also shown a significantly higher clutch size in earlier nests. As anthropogenic materials are higher in earlier nests, it is interesting to note that anthropogenic materials might be selected by higher-quality parents, especially in urban habitats.

In the study design moss and grass were weighed together considering them as natural materials. While moss was present in both the cup and base, the highest percentage was present in the base. On the other hand, the grass is seen only in the cup. Mostly anthropogenic material was used in cups instead of grass to provide structural integrity. Even though, the combination of combination of moss and grass showed a significant effect on breeding success it is important to find which item is contributing to this effect. Further studies are needed to confirm the effect specific to moss, grass and anthropogenic materials on breeding success. Also, the additional of availability of the materials in the surroundings can be used to assess the selective preference of nesting materials.

## Conclusion

Urbanization has a clear impact on the nesting materials chosen by birds. While anthropogenic materials are being used more frequently in nests, especially in urban environments, they do not necessarily predict breeding success as significantly as natural materials like moss and grass. This study emphasizes the importance of understanding how changing environments, and in particular urbanising ones, influence avian behaviour and reproductive fitness. Further research is essential to delve deeper into the specific roles of individual nesting materials and the choices birds make in different habitats.

## Acknowledgements

We thank Conor Haugh for the processing and collation of the environmental variables used in this study.

## Notes

### Competing Interest Statement

The authors have declared no competing interest.

### Summary of Updates

The previous version had editor comments which were unintended.

