## Supplementary Materia for "Urbanization Affects Nest Material Composition and Use of Anthropogenic Materials in Blue Tit Nests (*Cyanistes caeruleus)*" (from another version of this preprint: version 2, 2024-05-16)

Effect of Habitat and First Egg Date on Nest Materials

1) Total~ Habitat+ First Egg Date

| **Characteristic** | **Beta**^1^ | **SE**^2^ | **95% CI**^2^ | **p-value** |
| --- | --- | --- | --- | --- |
| (Intercept) | 38* | 15.8 | 5.8, 69 | 0.021 |
| first_egg_date | -0.15 | 0.131 | -0.42, 0.11 | 0.2 |
| Habitat |  |  |  |  |
| Rural | — | — | — |  |
| Urban | 1.9 | 2.73 | -4.6, 8.4 | 0.5 |
| ^1^*p<0.05; **p<0.01; ***p<0.001 | | | | |
| ^2^SE = Standard Error, CI = Confidence Interval | | | | |

2) Anthropogenic Material~ Habitat* First Egg Date

| **Characteristic** | **Beta**^1^ | **SE**^2^ | **95% CI**^2^ | **p-value** |
| --- | --- | --- | --- | --- |
| (Intercept) | -0.06 | 4.88 | -9.9, 9.7 | >0.9 |
| first_egg_date | 0.00 | 0.041 | -0.08, 0.08 | >0.9 |
| Habitat |  |  |  |  |
| Rural | — | — | — |  |
| Urban | 12 | 6.13 | -0.41, 24 | 0.058 |
| first_egg_date * Habitat |  |  |  |  |
| first_egg_date * Urban | -0.09 | 0.052 | -0.20, 0.01 | 0.077 |
| ^1^*p<0.05; **p<0.01; ***p<0.001 | | | | |
| ^2^SE = Standard Error, CI = Confidence Interval | | | | |

3) Moss~ Habitat+ First Egg Date

| **Characteristic** | **Beta**^1^ | **SE**^2^ | **95% CI**^2^ | **p-value** |
| --- | --- | --- | --- | --- |
| (Intercept) | 22 | 14.0 | -6.3, 50 | 0.13 |
| DATA$first_egg_date | -0.05 | 0.116 | -0.28, 0.18 | 0.7 |
| Habitat |  |  |  |  |
| Rural | — | — | — |  |
| Urban | -2.6 | 1.87 | -7.7, 2.5 | 0.2 |
| ^1^*p<0.05; **p<0.01; ***p<0.001 | | | | |
| ^2^SE = Standard Error, CI = Confidence Interval | | | | |

4) Feather ~ Habitat+ First Egg Date

| **Characteristic** | **Beta**^1^ | **SE**^2^ | **95% CI**^2^ | **p-value** |
| --- | --- | --- | --- | --- |
| (Intercept) | -0.21 | 3.07 | -6.4, 5.9 | >0.9 |
| DATA$first_egg_date | 0.01 | 0.025 | -0.04, 0.06 | 0.7 |
| Habitat |  |  |  |  |
| Rural | — | — | — |  |
| Urban | 1.0 | 0.974 | -1.7, 3.7 | 0.3 |
| ^1^*p<0.05; **p<0.01; ***p<0.001 | | | | |
| ^2^SE = Standard Error, CI = Confidence Interval | | | | |

5) Wool ~ Habitat+ First Egg Date

| **Characteristic** | **Beta**^1^ | **SE**^2^ | **95% CI**^2^ | **p-value** |
| --- | --- | --- | --- | --- |
| (Intercept) | 7.7 | 6.30 | -4.9, 20 | 0.2 |
| DATA$first_egg_date | -0.05 | 0.052 | -0.15, 0.06 | 0.4 |
| Habitat |  |  |  |  |
| Rural | — | — | — |  |
| Urban | 0.77 | 1.17 | -2.1, 3.7 | 0.5 |
| ^1^*p<0.05; **p<0.01; ***p<0.001 | | | | |
| ^2^SE = Standard Error, CI = Confidence Interval | | | | |

**Effect of IS100 and First Egg Date on Nest Materials**

6) Total~ IS100+ First Egg Date

| **Characteristic** | **Beta**^1^ | **SE**^2^ | **95% CI**^2^ | **p-value** |
| --- | --- | --- | --- | --- |
| (Intercept) | 21*** | 1.35 | 18, 24 | <0.001 |
| scale(first_egg_date) | -1.1 | 0.793 | -2.7, 0.48 | 0.2 |
| scale(IS100) | -1.5 | 1.06 | -3.7, 0.65 | 0.2 |
| ^1^*p<0.05; **p<0.01; ***p<0.001 | | | | |
| ^2^SE = Standard Error, CI = Confidence Interval | | | | |

7) Anthropogenic Materials ~ IS100+ First Egg Date

| **Characteristic** | **Beta**^1^ | **SE**^2^ | **95% CI**^2^ | **p-value** |
| --- | --- | --- | --- | --- |
| (Intercept) | 0.59 | 0.245 | -0.02, 1.2 | 0.056 |
| scale(first_egg_date) | -0.42* | 0.160 | -0.74, -0.09 | 0.012 |
| scale(IS100) | 0.20 | 0.201 | -0.23, 0.63 | 0.3 |
| ^1^*p<0.05; **p<0.01; ***p<0.001 | | | | |
| ^2^SE = Standard Error, CI = Confidence Interval | | | | |

8) Moss~ IS100+ First Egg Date

| **Characteristic** | **Beta**^1^ | **SE**^2^ | **95% CI**^2^ | **p-value** |
| --- | --- | --- | --- | --- |
| (Intercept) | 14*** | 0.635 | 13, 16 | <0.001 |
| scale(first_egg_date) | -0.38 | 0.668 | -1.7, 0.95 | 0.6 |
| scale(IS100) | -2.0** | 0.668 | -3.4, -0.69 | 0.004 |
| ^1^*p<0.05; **p<0.01; ***p<0.001 | | | | |
| ^2^SE = Standard Error, CI = Confidence Interval | | | | |

9) Feather~ IS100+ First Egg Date

| **Characteristic** | **Beta**^1^ | **SE**^2^ | **95% CI**^2^ | **p-value** |
| --- | --- | --- | --- | --- |
| (Intercept) | 1.5* | 0.500 | 0.08, 2.9 | 0.043 |
| scale(first_egg_date) | 0.04 | 0.161 | -0.28, 0.36 | 0.8 |
| scale(IS100) | -0.05 | 0.286 | -0.64, 0.54 | 0.9 |
| ^1^*p<0.05; **p<0.01; ***p<0.001 | | | | |
| ^2^SE = Standard Error, CI = Confidence Interval | | | | |

10) Wool and Hair ~ IS100+ First Egg Date

| **Characteristic** | **Beta**^1^ | **SE**^2^ | **95% CI**^2^ | **p-value** |
| --- | --- | --- | --- | --- |
| (Intercept) | 2.5** | 0.605 | 1.1, 3.9 | 0.004 |
| scale(first_egg_date) | -0.34 | 0.322 | -0.99, 0.30 | 0.3 |
| scale(IS100) | -0.15 | 0.452 | -1.1, 0.80 | 0.8 |
| ^1^*p<0.05; **p<0.01; ***p<0.001 | | | | |
| ^2^SE = Standard Error, CI = Confidence Interval | | | | |

**Effect of Nest Materials on Breeding Success (GLMM)**

11) ***(clutch_size ~ Anthro+Moss+Feather+ Wool.Hair+ (1|Site)+ first_egg_date, family = poisson)***

| **Characteristic** | **log(IRR)**^12^ | **SE**^2^ | **95% CI**^2^ | **p-value** |
| --- | --- | --- | --- | --- |
| (Intercept) | 2.9** | 0.998 | 0.98, 4.9 | 0.003 |
| Anthro | 0.02 | 0.037 | -0.05, 0.09 | 0.5 |
| Moss | 0.00 | 0.009 | -0.02, 0.02 | 0.7 |
| Feather | -0.01 | 0.043 | -0.10, 0.07 | 0.8 |
| Wool.Hair | 0.01 | 0.018 | -0.02, 0.05 | 0.5 |
| first_egg_date | -0.01 | 0.008 | -0.02, 0.01 | 0.4 |
| ^1^*p<0.05; **p<0.01; ***p<0.001 | | | | |
| ^2^IRR = Incidence Rate Ratio, SE = Standard Error, CI = Confidence Interval | | | | |

12) ***(hatch_per ~ Anthro + Moss + Feather + Wool.Hair + (1|Site)+ first_egg_date, family = binomial)***

| **Characteristic** | **log(OR)**^12^ | **SE**^2^ | **95% CI**^2^ | **p-value** |
| --- | --- | --- | --- | --- |
| (Intercept) | -9.2 | 20.2 | -49, 30 | 0.6 |
| Anthro | 0.40 | 1.23 | -2.0, 2.8 | 0.7 |
| Moss | 0.09 | 0.277 | -0.45, 0.64 | 0.7 |
| Feather | -0.31 | 0.917 | -2.1, 1.5 | 0.7 |
| Wool.Hair | -0.26 | 0.381 | -1.0, 0.49 | 0.5 |
| first_egg_date | 0.11 | 0.171 | -0.22, 0.45 | 0.5 |
| ^1^*p<0.05; **p<0.01; ***p<0.001 | | | | |
| ^2^OR = Odds Ratio, SE = Standard Error, CI = Confidence Interval | | | | |

13) *(fledg_per ~ Moss + (1|Site)+ first_egg_date, family = binomial)*

| **Characteristic** | **log(OR)**^12^ | **SE**^2^ | **95% CI**^2^ | **p-value** |
| --- | --- | --- | --- | --- |
| (Intercept) | -8.2 | 9.35 | -26, 10 | 0.4 |
| Moss | 0.27 | 0.147 | -0.02, 0.55 | 0.069 |
| first_egg_date | 0.06 | 0.077 | -0.09, 0.21 | 0.4 |
| ^1^*p<0.05; **p<0.01; ***p<0.001 | | | | |
| ^2^OR = Odds Ratio, SE = Standard Error, CI = Confidence Interval | | | | |

14) **Model Selection for Fledling Success**

*Model1 <- glmer(fledg_per ~ Moss + (1|Site)+ first_egg_date,*

*data = DATA, family = binomial)*

*Model2 <- glmer(fledg_per ~ Anthro + Moss + Feather + Wool.Hair + (1|Site)+*

*first_egg_date,data = DATA, family = binomial)*

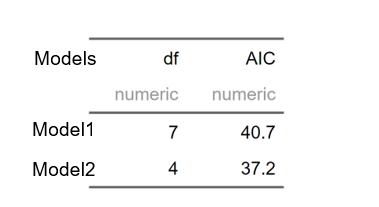
